# A modular T7-based gene expression platform in *Pseudomonas putida*: Construction and *in silico* analysis

**DOI:** 10.1101/2022.06.02.494349

**Authors:** Marleen Beentjes, Hannes Löwe, Katharina Pflüger-Grau, Andreas Kremling

## Abstract

The T7 RNA polymerase is considered one of the most popular tools for heterologous gene expression in the gold standard biotechnological host *Escherichia coli*. However, the exploitation of this tool in other prospective biotechnological hosts is still very scarce. To this intent, we established and characterized a modular T7 RNA polymerase-based system for heterologous protein production in *Pseudomonas putida*, using the model protein eGFP as an easy-to-quantify reporter protein. We have effectively targeted the limitations associated with the initial genetic set-up of the system, such as slow growth and low protein production rates. By replacing the T7-phage inherent TΦ terminator downstream of the heterologous gene with the synthetic tZ terminator, growth and protein production rates improved drastically, and the T7 RNA polymerase system reached a productivity level comparable to that of an intrinsic RNA polymerase based system. Furthermore, we could show that the system is saturated with T7 RNA polymerase by applying a T7 RNA polymerase ribosome binding site library to tune heterologous protein production. The saturation points to an essential role for the ribosome binding sites of the T7 RNA polymerase since, in an oversaturated system, cellular resources are lost to the synthesis of unnecessary T7 RNA polymerase. Eventually, we combined the experimental data into a model that can predict the eGFP production rate with respect to the relative strength of the ribosome binding sites upstream of the T7 gene.

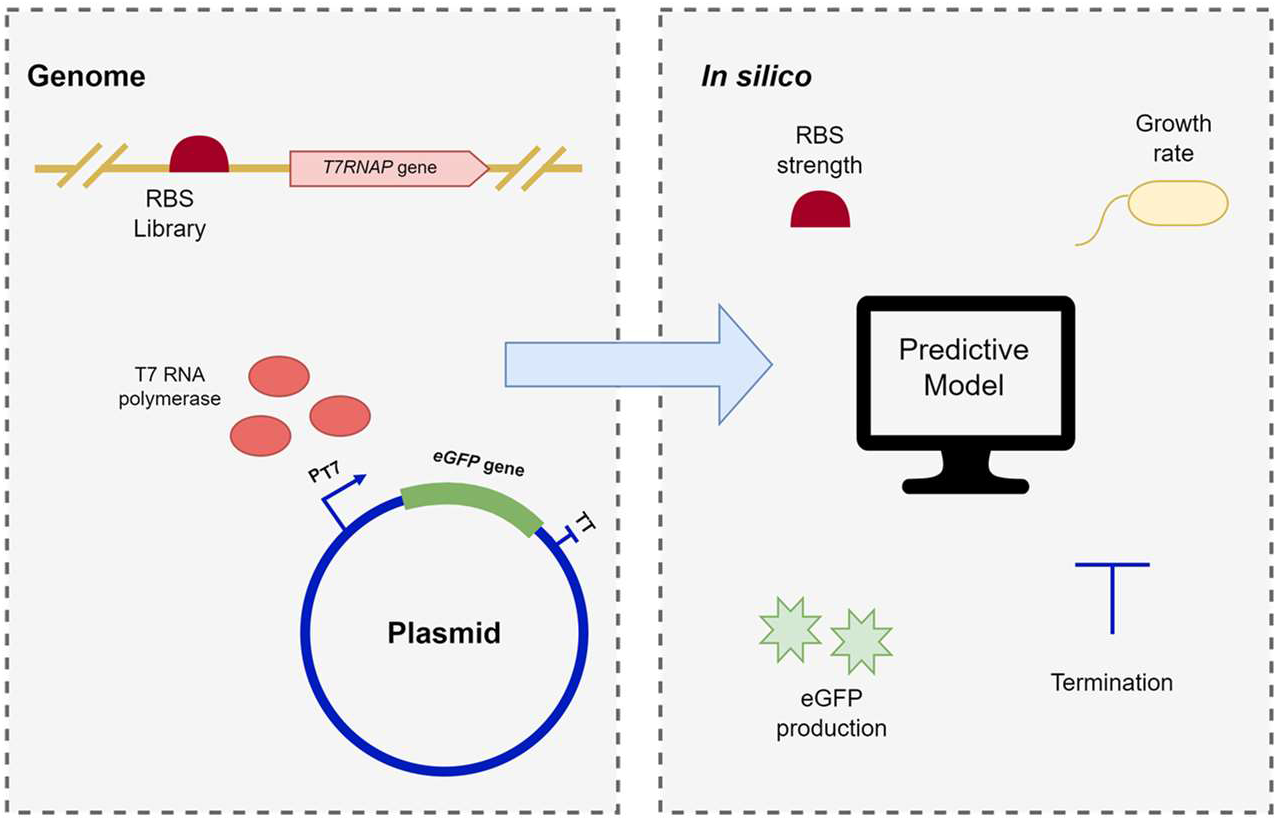

## Introduction

The gram-negative bacterium *Pseudomonas putida* has emerged as a potential host for various biotechnological applications during the past decades.^*1*^ The bacterium’s natural presence in different environments provides it with widespread chemo- and physical tolerance and high adaptability skills.^*2*^ Furthermore, *P. putida* is characterized by fast growth and the ability to utilize various carbon sources.^*3*^ Driven by the advantageous natural traits of *P. putida*, efforts were made to further expand its potential and a large toolbox for the genetic enhancement of *P. putida* has been developed until now. This toolbox consists of classical methods, like the Tn5 and Tn7 mini plasmid-based genomic insertion, as well as newer CRISPR-Cas9-based recombination methods.^*4*^ In addition, the compatibility of *P. putida* with plasmids provided by the Standard European Vector Architecture (SEVA) platform significantly reduces the effort for the construction of various recombinant *P. putida* strains.^*5*^

With the availability of these genetic tools, the exploitation of *P. putida* as a host for heterologous gene expression increased.^*6-8*^ Heterologous gene expression has been well studied in recombinant *Escherichia coli*, which is one of the most popular production hosts in research. More specifically, many established *E. coli* strains for heterologous gene expression rely on the T7 RNA polymerase (T7RNAP) for the transcription of desired genes.^*9*^ The T7RNAP is an RNA polymerase, originating from the T7 bacteriophage, with a very high affinity for its associated T7 promoter. Cloning a gene of interest (GOI) downstream of the T7 promoter leads to high-level, specific, and constitutive expression of this gene when T7RNAP is present in the system.^*10*^ Despite the vast amount of research on T7RNAP-based expression systems in *E. coli*, research on T7RNAP-based systems in *P. putida* is limited.^*11-14*^

Different genetic constructs using the T7RNAP in *P. putida* have been described in the past: One is the expression of the T7RNAP from a plasmid. Arvani et al. constructed the vector pML5-P_lac_T7, harboring the T7RNAP gene under the control of the IPTG inducible *lacUV5* promoter.^*11*^ After conjugation of this plasmid into *P. putida* KT2440, the resulting strain successfully produced T7RNAP upon induction with IPTG.

Another plasmid-based T7RNAP system was designed by Liang et al.^*12*^ In their host independent T7 expression system (HITES), the T7RNAP gene and GOI are both located on the same plasmid. Several genetic elements were introduced or adapted to fine-tune the expression. The HITES enabled GFP production in various gram-negative bacterial strains, including *P. putida*. However, since already small amounts of T7RNAP suffice for high-level production of a GOI under the control of the T7 promoter, a higher copy number of the T7RNAP gene on a plasmid will not be very beneficial.^*15*^

Examples where the T7RNAP gene was introduced into the chromosome of *P. putida* also exist. The first to do this were Herrero et al.: In their system, both, T7RNAP, under the control of the 3-methyl benzoate (3-MB) inducible XylS/*Pm* promoter, and a recombinant gene (*LacZ*) were randomly integrated into the genome of *P. putida* using a mini Tn5-plasmid.^*16*^ To evaluate the performance of their system, the product of the T7RNAP driven *LacZ* expression, β-Galactosidase, was quantified in *P. putida* and *E. coli*. Upon induction with 3-MB, both strains produced nearly the same amount of β- Galactosidase. Without 3-MB induction, no β-Galactosidase production could be observed for *P. putida*, whereas for *E. coli*, substantial β-Galactosidase production was observed also in the absence of 3-MB.

Troeschel et al. inserted the T7RNAP gene under the control of the IPTG inducible *lacUV5* promoter into the genome of *P. putida* KT2440.^*13*^ They used a mini-Tn7 plasmid for specific integration into the *att*Tn7 side. Plasmids containing lipase A from *Bacillus subtilis* or cutinase from *Fusarium solanipisi* were used to verify heterologous protein production. The activity of the enzymes in the supernatant and cell pellet of the *P. putida* KT2440-T7 strain was distinctively lower than in the T7RNAP-producing *E. coli* (BL21) reference strain. For both strains, background enzyme activity was found in the absence of the T7RNAP inducer IPTG.

Calero et al. characterized and compared a range of commonly applied promoters for the production of heterologous proteins in *P. putida*.^*14*^ Analogous to Troeschel et al., they constructed a *P. putida* KT2440 strain harboring the T7RNAP gene in the genome, also applying the *lacUV5* promoter to mediate the expression of the T7RNAP. A plasmid with GFP under the control of the LacI/*T7* promoter was conjugated into that same strain. The performance of this strain was compared to wild-type KT2440 strains with the same GFP expression plasmid, but with different (non T7RNAP dependent) promoters (*ParaB, PlacUV5*, XylS*/Pm, Psal, and PalkB)*. The strain with the XylS*/Pm* promoter performed best with respect to the heterologous protein expression, and the strains with the LacI*/T7* and the *PlacUV5* showed the lowest expression levels. These three promoters (XylS*/Pm*, LacI*/T7* and *PlacUV5*) also caused the highest level of basal expression in comparison to the other tested promoters. Moreover, a decrease in the growth rate of *P. putida* was observed for the LacI*/T7* and XylS*/Pm* strains.

The studies discussed in the previous paragraphs indicate the applicability of different T7RNAP-dependent expression systems in *P. putida*. However, there is a lack of explanation as to why certain genetic set-ups show low expression strengths and possess only weak controllability. This basic understanding of the systems is important considering the need for predictable and robust biotechnological processes. In this work, we established and thoroughly characterized a T7RNAP-dependent system for the production of heterologous proteins in *P. putida*. Its performance was compared to a well-established T7RNAP-independent expression system. Furthermore, we engineered suitable strains to differentially monitor the influence of transcriptional and translational processes on the overall activity of the T7RNAP-dependent expression system. To get a deeper understanding of the system, we separately inspected the influence of transcription termination on the system by exchanging the terminator downstream of the GOI. Moreover, a library of ribosome binding sites for T7RNAP expression was created to analyze its role in tuning the expression of the GOI. The experimental data were applied to set up a mathematical model, that was able to predict the relative strength of a ribosome binding site from the eGFP production rate. The information derived from the model can be used to suggest the genetic set-up that gives the best results concerning productivity and controllability.

## Results and Discussion

### Construction and characterization of the T7RNAP based expression system

In the T7RNAP expression system described here, the *T7RNAP* gene was integrated into the chromosome under the control of the XylS/*Pm* promoter, and the gene encoding the reporter protein, the enhanced Green Fluorescent Protein (eGFP), was located on a pSEVA derived plasmid (Figure 1). Since the *T7RNAP* and *eGFP* genes are situated in separated genetic locations, the system is considered to be modular. This kind of system poses an advantage over non-modular systems because both host and plasmids can be freely combined to tune the expression of the GOI.

**Figure 1.**
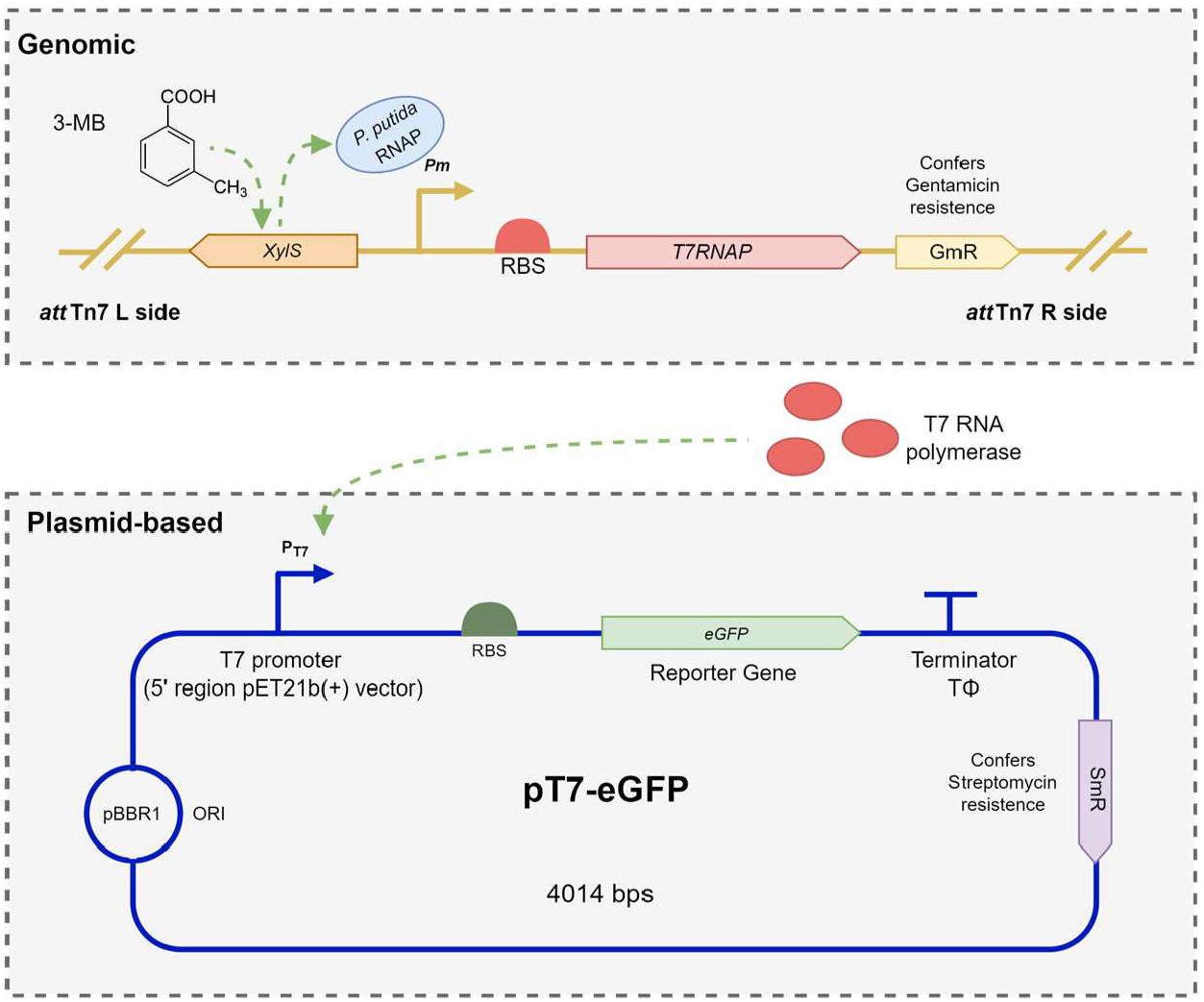
Genetic layout of the T7RNAP-based expression system. The *T7RNAP* is integrated into the *att*Tn7 side of the genome of *P. putida* KT2440 and its expression is under the control of the 3-MB inducible XylS/*Pm* promoter. The T7RNAP in turn initiates *eGFP* transcription from the T7 promoter (originating from the 5’ region upstream of the promoter of the pET21b(+) vector). The T7 bacteriophage’s inherent TΦ terminator is located downstream of the *eGFP* gene. The pT7-eGFP vector confers streptomycin resistance (SmR) to the plasmid harboring cells.

To create the first module of the system, a mini pTn7 plasmid, designated to deliver the T7RNAP gene into the *att*Tn7 side of the *P. putida* genome, was constructed. The pTn7-XylS/*Pm*-T7 plasmid was conjugated into *P. putida* KT2440, giving rise to *P. putida* KT2440 *att*Tn7::XylS/*Pm*-T7, which from now on for the sake of clarity will be referred to as *P. putida* T7RNAP. The second module of the system compromises a T7 promotor driven expression plasmid, called pT7-eGFP. This plasmid was constructed using the standard pSEVA438 plasmid as a template. The promoter cassette XylS/*Pm* of the pSEVA438 plasmid was exchanged with the 5’ region of the BL21 expression plasmid pET21b(+), containing the T7RNAP specific T7 promoter. The reporter construct (composed of a spacer optimized RBS^*17*^, *eGFP* and the T7 phage inherent TΦ terminator) was introduced downstream of the T7 promoter. The plasmid pT7-eGFP was then introduced into the aforementioned *P. putida* T7RNAP strain by conjugation. In the resulting strain, *P. putida* T7RNAP (pT7-eGFP), T7RNAP is produced upon induction of the XylS/*Pm* promoter system with 3-MB. The T7RNAP in turn recognizes the T7 promoter on the plasmid and transcribes the *eGFP*, which is located downstream of the promoter. The fluorescence of eGFP can in turn be measured to evaluate the performance of the system. Both the mini Tn7-based genomic T7RNAP cassette and the pSEVA-derived expression plasmid are compatible with a variety of other gram-negative hosts.

Before the discovery of orthogonal expression systems like the T7-system, heterologous protein production relied solely on the host’s intrinsic, non-specific RNA polymerases for the transcription of the target recombinant gene. To obtain a reference for the performance of the T7RNAP-based system in *P. putida*, we compared it to a well-established expression system in *P. putida*, which only relies on the native machinery for heterologous protein production (see Figure 2). Therefore, *P. putida* KT2440 was equipped with a pSEVA438-eGFP vector, expressing *eGFP* upon induction of the XylS/*Pm* promoter with 3-MB. In this strain, *eGFP* transcription is accomplished by the host’s natural RNA polymerases. In contrast to the T7RNAP, which is dedicated exclusively to the transcription of genes downstream of the T7 promoter, inherent RNA polymerases are also responsible for native gene transcription in the host. Therefore, T7RNAP is considered to be more efficient for heterologous gene expression.

**Figure 2.**
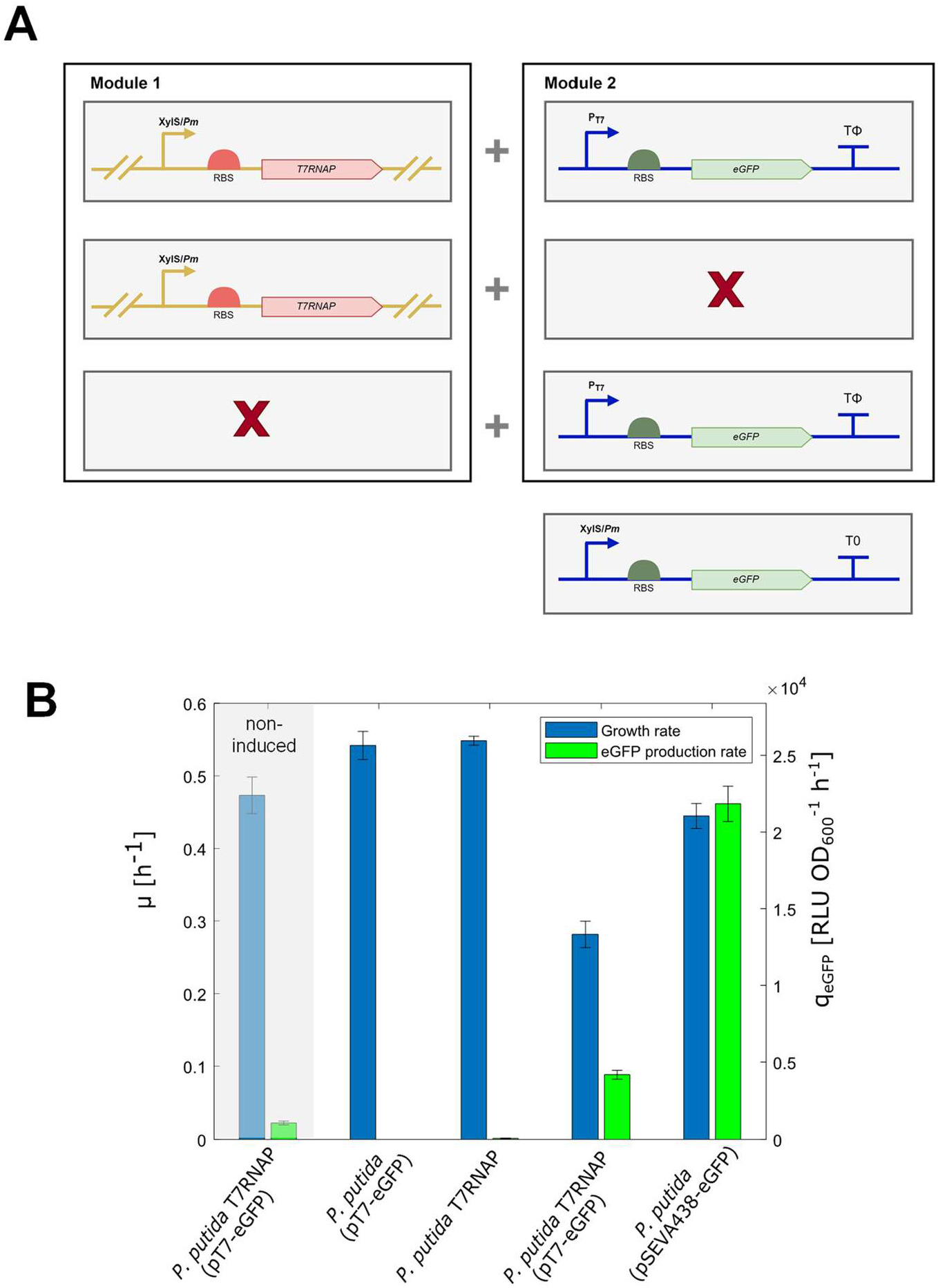
Verification of the T7RNAP expression system in *P. putida* KT2440. **A:** Genetic layout of the strains that were applied to test the system. Module 1 comprises the *T7RNAP* gene integrated into the genome and module 2 a plasmid for T7RNAP driven *eGFP* expression. In total, four strains were tested; one strain that contained both module 1 and module 2, two strains that contained either module 1 or module 2, respectively, and one reference strain that contained an adapted second module (plasmid relying on host intrinsic RNAP for transcription). **B:** Growth rate and eGFP production rate. Cultures were grown in 10 mL M9 Minimal medium with 3 g L^-1^ glucose as a C-source, at a temperature of 30°C and an agitation speed of 220 rpm. Induction of the cultures was achieved by adding 0.1 mM 3-MB at the start of cultivation. The grey shaded area represents the results of the non-induced *P. putida* T7RNAP (pT7-eGFP) as control. The cultivations were performed in triplicates for all strains. Error bars represent the standard deviation (based on a sample) of the replicates, calculated using the n-1 method.

When analyzing the strains *P. putida* T7RNAP and *P. putida* (pT7-eGFP), which both carry only one of the two modules, no eGFP production could be detected. Only the combined expression of T7RNAP and presence of the pT7-eGFP plasmid resulted in detectable eGFP fluorescence, confirming the orthogonality of the system. However, the specific eGFP production rate (q_eGFP_) in the modular T7-based system was 4-fold lower than observed for the host intrinsic RNA polymerase-based system in *P. putida* (pSEVA438-eGFP). This was unexpected since the specificity of the T7RNAP for the T7 promoter should theoretically allow the system to produce more protein than the reference system. Moreover, the growth rate (µ) of *P. putida* T7RNAP (pT7-eGFP) was decreased by almost a factor of 2 in comparison to the strains with only one modular part. This suggests that a certain type of stress is imposed on the strain when eGFP is produced. Furthermore, the *P. putida* T7RNAP (pT7-eGFP) was able to produce eGFP even in the absence of 3-MB, indicating that the XylS/*Pm* is not entirely tightly regulated in this system’s set-up, allowing for a certain level of T7RNAP basal expression. The controllability of the system, defined by dividing the eGFP production rate in an induced state by the production rate in a non-induced state, was only a factor of 3.9. As the tunability of a system is an important asset in biotechnological processes, a higher controllability factor would be desirable. The following experiments, therefore, were aimed to identify the nature of the stress that caused the decreased growth and the poor controllability.

### Discerning transcriptional and translational effects

The first strategy was to verify whether the stress caused by combined T7RNAP and eGFP production in the modular system originated from transcriptional or translational problems. Therefore, two additional strains were constructed. Both strains are essentially identical to *P. putida* T7RNAP (pT7-eGFP), but carry variations in the RBS upstream of the *eGFP* gene. One strain, *P. putida* T7RNAP (pT7-eGFP-ΔRBS), does lacks the RBS upstream of the *eGFP* gene, disabling translation of the eGFP messenger RNAs. Hence, this modification should only influence the transcriptional capacity of the strain. The other strain, *P. putida* T7RNAP (pT7-eGFP-strongRBS), contains a stronger RBS designed by Ceroni et al.^*18*^ This synthetic RBS was optimized for efficient translation of superfolding GFP by using the Ribosome Binding Site Calculator.^*19*^ Accordingly, the presence of this stronger RBS upstream of the eGFP mRNA transcript is expected to pose an increased pressure on the translation machinery of the cells.

As suggested, *P. putida* T7RNAP (pT7-eGFP-strongRBS) with the strong RBS upstream of the *eGFP* gene, had the highest rate of eGFP production (Figure 3). However, this increase in eGFP production rate, and therefore enhanced pressure on the translation machinery of the strain, was not accompanied by a visible decrease in growth rate compared to that of *P. putida* T7RNAP (pT7-eGFP). The *P. putida* T7RNAP (pT7-eGFP-ΔRBS) strain, which lacks the capacity to translate the eGFP mRNA into proteins, still suffered from a reduced growth rate compared to the non-induced control. This indicates that the cause of reduced growth in the T7RNAP strains does not lie at the translational level, but rather has a transcriptional origin. This phenomenon of transcriptional burden in a T7RNAP system was also already observed for *E. coli* (BL21) expressing recombinant proteins.^*20*^

**Figure 3.**
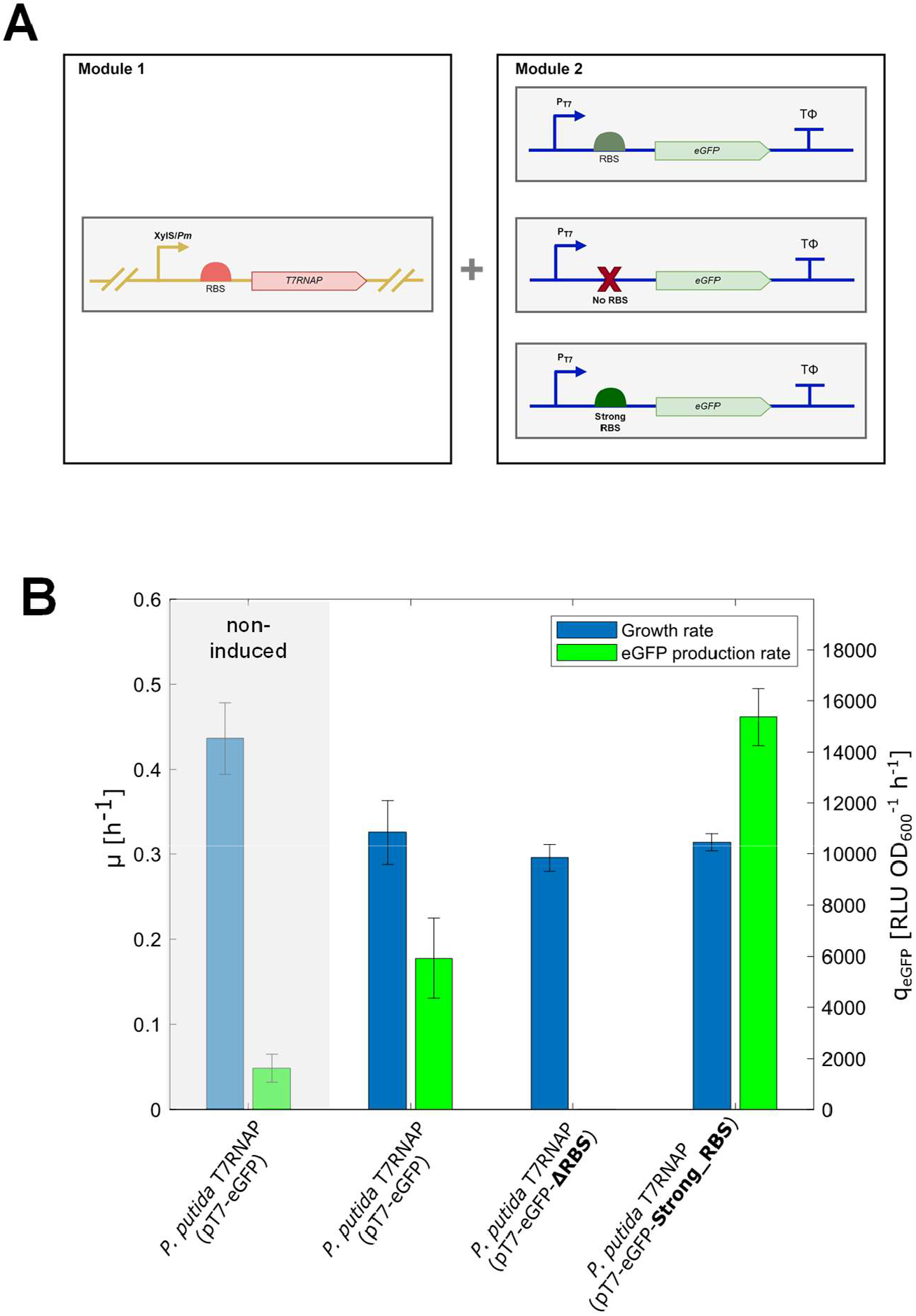
Discerning transcriptional and translation problems in the T7RNAP expression system. **A:** Genetic layout of the strains that were applied to test the system. Module 1 comprises the *T7RNAP* gene integrated into the genome, module 2 comprises a plasmid for T7RNAP driven *eGFP* expression. In total, three strains were tested; Module 1 was the same for all three strains, but the composition of module 2 varied in the RBS. **B:** Growth rate and eGFP production rate. Cultures were grown in 10 mL M9 Minimal medium with 3 g L^-1^ glucose as a C-source, at a temperature of 30°C and an agitation speed of 220 rpm. Induction of the cultures was achieved by adding 0.1 mM 3-MB at the start of cultivation. The grey shaded area represents the results of non-induced *P. putida* T7RNAP (pT7-eGFP) as control. The cultivations were performed in triplicates for all strains. Error bars represent the standard deviation (based on a sample) of the replicates, calculated using the n-1 method.

### Influence of the terminator on transcription

To come up with a strategy to mitigate the effects of this transcriptional burden, we set out to investigate the role of the T7 bacteriophage inherent TΦ terminator, downstream of the *eGFP* gene, in the transcription process. Previous research already indicated that the application of this specific terminator is prone to cause ‘read-through’ transcription.^*21-23*^ This occurs when the terminator sequence fails to abort the transcription process of an RNA polymerase, causing a consecutive transcription downstream of the terminator sequence. Research by Dunn and Studier has suggested that for the T7 bacteriophage, this read-through is necessary since the genes downstream of the TΦ terminator do not contain additional promotor sequences, even though they are essential for the reproduction of the bacteriophage.^*24*^ This means that the transcription of these genes relies solely on the read-through of the T7RNAP. For the transcription of a gene on a synthetic plasmid, this effect however is not desirable and can cause several problems for the cells, e.g. a loss of cellular resources to the synthesis of unnecessary long mRNAs or the disturbance in plasmid replication due to read-through into the origin of replication.

To test this hypothesis, we performed an *in vitro* transcription assay to assess whether a read-through downstream of the *eGFP* could be observed (Figure 4A). Before the assay, we generated transcription templates of different lengths by performing a PCR using pT7-eGFP as a template. All transcription templates contained the T7 promoter sequence, the *eGFP* coding sequence and the TΦ terminator, but they differed in their sequence length downstream of the TΦ terminator. The reverse primers that were used for this PCR were named “ReadThrough” 1 to 5, with each consecutive number indicating that the resulting PCR fragment contained ca. 500 base pairs more of the plasmid sequence downstream of the TΦ terminator than the previous one. The standard fragment contained just the T7 promoter sequence, the *eGFP* coding sequence and the TΦ terminator. The *in vitro* transcripts of the longer DNA fragments should match the length of the transcript of the standard fragment, if the transcription is correctly aborted at the terminator sequence. Separate T7 RNA polymerase transcription assays were performed with the differently sized PCR fragments as templates, as well as a negative control assay without T7RNAP. This control was also to verify that the DNA template used for the transcription assay is not visible on the RNA agarose gels. As can be seen in Figure 4B, the resulting transcripts of the different assays did not have the same size as the transcript “TΦ” derived from the reference fragment, indicating that the TΦ terminator did not effectively prevent read-through transcription in an *in vitro* system.

**Figure 4.**
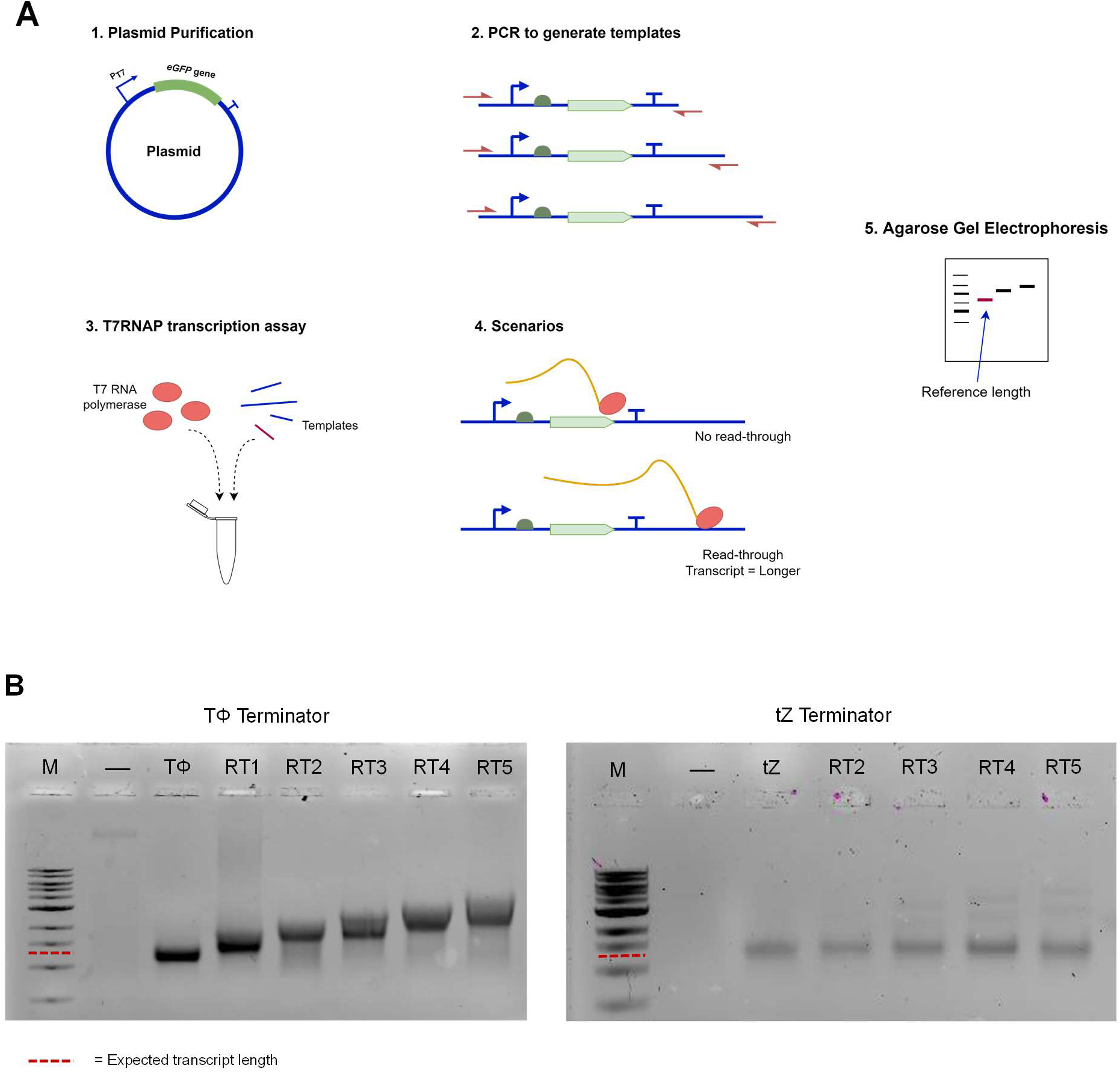
Workflow and results of the T7RNAP *in vitro* transcription assay for verification of read-through transcription by the T7RNAP. **A:** Graphical representation of the different steps for *in vitro* assessment of T7RNAP read-through. Shortly, a PCR is performed using the purified plasmids pT7-eGFP or pT7-eGFP-tZ as a template. By varying the reverse primer of the PCR reactions, fragments with the T7 promotor, *eGFP* gene, terminator and a variable length of sequence downstream of the terminator were generated. These fragments were used as a template in the subsequent *in vitro* transcription assays. The length of the transcripts for each assay was then determined by gel electrophoresis. **B:** RNA Agarose gel showing the transcripts of the T7 RNA polymerase transcription assays with differently sized DNA templates (left panel: containing the TΦ terminator sequence; right panel: containing the tZ terminator sequence). Lanes: M: 1 kb-DNA ladder (only for reference, not for estimation of transcript length), “- “ indicates the negative control, an assay without T7RNAP. TΦ / tZ indicates the transcripts of the DNA template that encompasses only the T7 promoter, *eGFP* gene and a terminator. RT1 to 5 indicate the transcripts of the longer “read-through DNA templates” that were used for the transcription assay. The dotted red line marks the expected transcript length for the case where no read-through occurs.

To test whether the observed *in vitro* read-through was contributing to the slow growth and low eGFP production rate associated with the expression system, we created a new strain, *P. putida* T7RNAP (pT7-eGFP-tZ). In this strain, the TΦ Terminator was exchanged with the tZ terminator taken from Mairhofer et al.^*21*^ The tZ terminator is composed of three different naturally occurring terminators: the class I T3TΦ terminator, the T1 termination signal of the *rrnB* gene and the TΦ terminator. The tZ terminator has a reported termination efficiency (TE) of 99%.^*21*^ For comparison, the wild-type TΦ terminator has a TE of approximately 79%. The tZ terminator has already been successfully applied to prevent read-through by several research groups.^*25-28*^ We used the newly constructed pT7-eGFP-tZ plasmid to generate DNA fragments for a new set of T7 RNA polymerase transcription assays in the same fashion as described above for the pT7-eGFP plasmid with the TΦ terminator. In this case, however, the DNA fragment “tZ” (containing the T7 promoter, *eGFP* and the tZ terminator only), is the fragment that after transcription yields the reference length mRNA transcript. From Figure 4B, it becomes apparent that, in contrast to the assay with the TΦ terminator, all transcription assays with the tZ terminator yielded RNA transcripts of about the same length as that of the tZ reference fragment. For the assays with DNA templates 3 to 5 however, some weak bands could be observed, which point to the presence of some larger transcripts. This is to be expected since even though the tZ terminator has a distinctly higher TE than its TΦ counterpart, the TE does still not amount to 100%. Nevertheless, these results indicate that the tZ successfully prevented read-through in an *in vitro* environment.

### Stronger transcription termination leads to a reduction of burden

The next step was to assess the effect of improved TE on the growth and eGFP production of the *P. putida* expression platform strain. The replacement of the TΦ terminator with the synthetic tZ terminator positively affected both aspects, raising the growth rate by roughly 25% and the eGFP production rate by almost 250% (Figure 5). Different hypotheses could explain the superior performance of the tZ system compared to the original system. As discussed in the previous section, read-through can potentially lead to the disruption of plasmid replication or a loss of resources in the synthesis of unwanted mRNA. By replacing the TΦ terminator and increasing the TE, these effects could be mitigated to some extent. This promotes overall improved cell viability and the availability of additional resources that would otherwise have been lost in read-through transcription.

**Figure 5.**
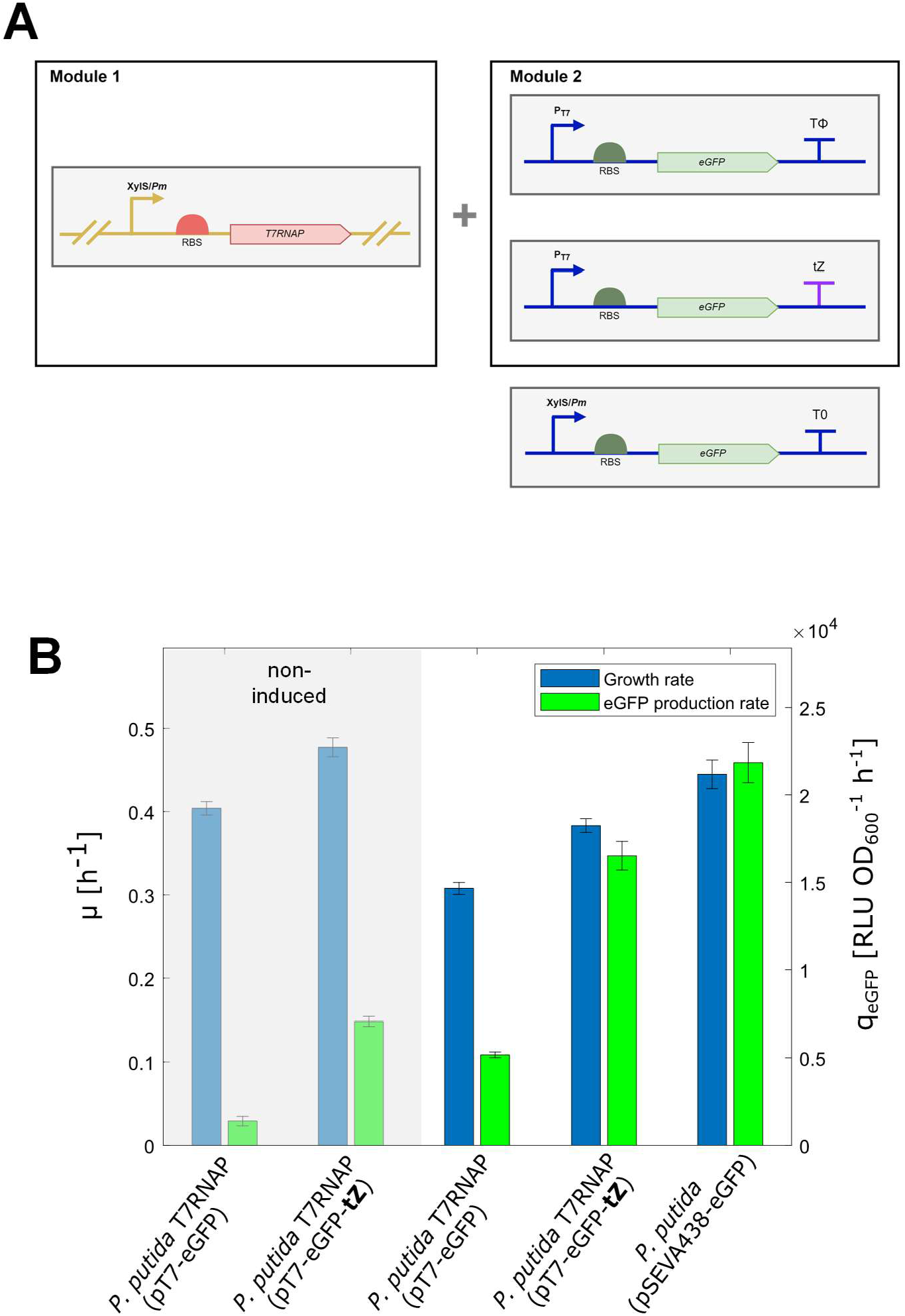
Effect of improved TE on growth rate and eGFP production for *P. putida* T7RNAP (pT7-eGFP-tZ). **A:** Genetic layout of the strains that were applied to test the effect of improved TE. Module 1 comprises the *T7RNAP* gene integrated into the genome, module 2 comprises a plasmid for T7RNAP driven *eGFP* expression. In total, three strains were tested; one strain that contained both, module 1 and module 2, one strain where the TΦ terminator of module 2 was exchanged with the tZ terminator and one reference strain that only contained an adapted second module (plasmid relying on host intrinsic RNAP for transcription). **B:** Growth rate and eGFP production rate. Cultures were grown in 10 mL M9 Minimal medium with 3 g L^-1^ glucose as a C-source, at a temperature of 30°C and an agitation speed of 220 rpm. Induction of the cultures was achieved by adding 0.1 mM 3-MB at the start of cultivation. The grey shaded area represents the results of the non-induced strains as control. All cultivations were performed in triplicates. Error bars represent the standard deviation (based on a sample) of the replicates, calculated using the n-1 method.

Coming back to the previously discussed work on T7RNAP systems in *P. putida*, our findings can explain why some of these expression systems show weak expression strengths and expression-related growth problems. The system tested by Troeschel et al. relied on an adaptation of the conventional pWE15 plasmid for transcription of their GOIs by the T7RNAP.^*13*^ This plasmid does not possess any terminator specific for the T7RNAP downstream of the GOI, making read-through transcription by the T7RNAP a likely occurrence. This could explain the low expression level of the GOIs observed in *P. putida*. Calero et al. applied the plasmid pRSFDuet-1 for their T7-based expression system.^*14*^ In their comparison of expression systems, the T7 system showed the weakest expression levels and was also reported to suffer from slow growth. These observations might also be explained by the presence of the T7 inherent TΦ terminator downstream of the GOI in the pRSFDuet-1 plasmid.

However, two side notes have to be added regarding the effectiveness of the system described here. Even though the system with the tZ terminator displayed a distinctively better performance than the previous version of the system, the T7RNAP-based tZ system did not outperform the reference host-intrinsic RNA polymerase system. The eGFP production rate and growth rate were slightly lower in the tZ system in comparison to the pSEVA system. The controllability, i.e. the eGFP production rate in the induced state divided by the rate in the non-induced state, was slightly lower for *P. putida* T7RNAP (pT7-eGFP-tZ) compared to *P. putida* T7RNAP (pT7-eGFP) (2.3 vs 3.7, respectively). Therefore, we came up with a strategy that would allow us to tune and closely inspect the system by varying the amount of T7RNAP.

### Testing the T7RNAP saturation hypothesis: construction of an RBS library

Another hypothesis for the poor performance of the original T7RNAP system is the potential saturation of transcription with T7RNAP. Since the T7RNAP, unlike the host-intrinsic polymerases, is dedicated exclusively to the transcription of the genes downstream of the T7 promoter, “small” amounts of the polymerase would theoretically suffice for the efficient transcription of the genes of interest. Once the system is saturated with T7RNAP, an increase in T7RNAP should no longer significantly enhance the eGFP production rate, meaning that unnecessary cellular resources will flow into the synthesis of non-useful T7RNAP. This saturation effect can also be described mathematically, where the rate of transcription is described by a function that contains a saturation term involving polymerase concentration.^*29*^ To test our hypothesis, we constructed a T7RNAP ribosome binding site library to govern the production of T7RNAP. This library included four degenerated variations of the original T7RNAP ribosome binding site (Table 1). In addition to this, an RBS from literature^*30*^ (defined as a medium-strength RBS for the production of yellow fluorescent protein) was used to create a reference to previous works on RBS strength. The expected expression level of the *T7RNAP* gene downstream of the different RBS in the library was identified using the UTR-designer tool from Seo et al.^*31*^ All experiments with the RBS library were initially performed with strains containing the TΦ terminator downstream of the *eGFP* gene. The results of these experiments are shown in Figure 6.

**Table 1.** Ribosome binding sites, conserved sequences, and accompanying expression values predicted by the UTR designer. Degenerated bases are depicted in red.

| RBS name | Conserved RBS Sequence | UTR Expression value |
| --- | --- | --- |
| Original | GGAAGAGG | 792575.07 |
| Deg 1 | GGAAGAGA | 143943.70 |
| Deg 2 | GTAAGAGA | 100070.73 |
| Deg 3 | GTAATAGA | 20325.42 |
| Deg 4 | GTAATCGA | 6021.85 |
| Medium | TTCGCAGGG | 20325.42 |

**Table 2.** Estimated parameter values k and K for the RBS library of *P. putida* T7RNAP (pT7-eGFP)

| Strain | condition / parameter | $k$ [a.u.] | $K$ [a.u.] |
| --- | --- | --- | --- |
| *P. putida* T7RNAP (pT7-eGFP) | standard induced | 5.359 | $4.29 \cdot 10^3$ |
| *P. putida* T7RNAP (pT7-eGFP) | standard non-induced | 1.398 | $4.29 \cdot 10^3$ |

**Table 3.** Alteration of parameters *k* and *K* by factors *α* and *β*, respectively, upon changing either termination of eGFP transcription or translation initiation of the eGFP transcript.

| Strain | condition / parameter | $\alpha$ [-] | $\beta$ [-] |
| --- | --- | --- | --- |
| *P. putida* T7RNAP (pT7-eGFP-tZ) | modified termination induced<br>(compared to standard induced) | 4.30 | 1.43 |
| *P. putida* T7RNAP (pT7-eGFP-tZ) | modified termination non-induced<br>(compared to standard non-induced) | 4.30 | 1.43 |
| *P. putida* T7RNAP (pT7-eGFP-strongRBS) | Strong RBS upstream of *eGFP* gene<br>(compared to standard induced) | 2.86 | 0.60 |

**Figure 6.**
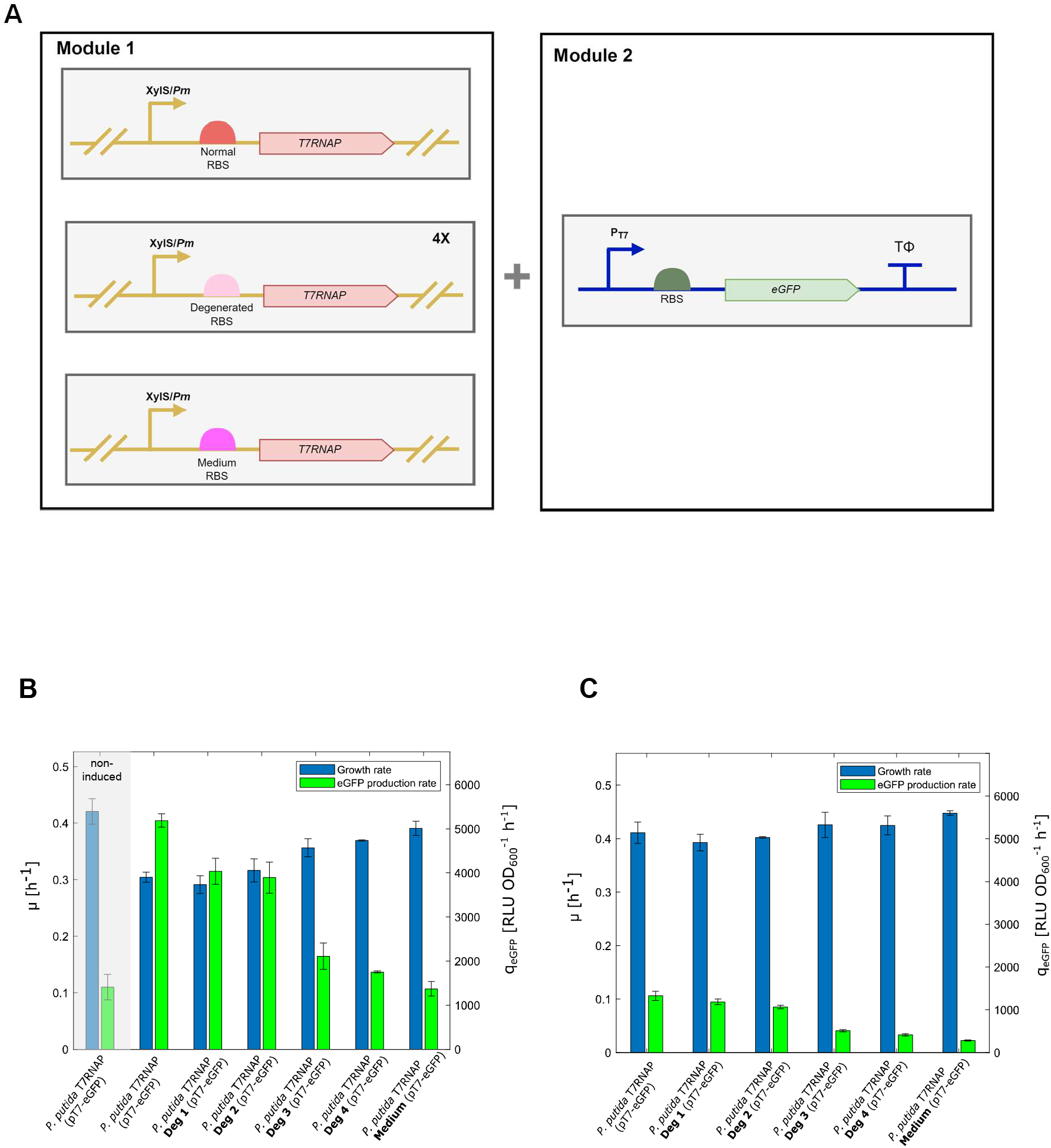
Tuning of eGFP production by applying a T7RNAP RBS library. **A:** Genetic layout of the strains that form the RBS library. Module 1 comprises the *T7RNAP* gene integrated into the genome, module 2 comprises a plasmid for T7RNAP driven eGFP expression. In total, six strains were tested. The difference between the strains was the RBS upstream of the *T7RNAP* gene in module 1. The predicted strength of the RBS sequences is gradually decreasing (compare table 1). Module 2 had the same composition for all strains. **B + C:** Growth rate and eGFP production rate. Cultures were grown in 10 mL M9 Minimal medium with 3 g L-1 glucose as a C-source, at a temperature of 30°C and an agitation speed of 220 rpm. The cultures were cultivated either with 0.1 mM 3-MB (**B**) or without any inducer (**C**). The grey shaded area in B represents the results of the non-induced strains as control. The cultivations were performed in duplicates for all strains. Error bars represent the standard deviation (based on a sample) of the replicates, calculated using the n-1 method.

Even though the UTR designer predicted a 5-fold higher T7RNAP expression level with the original RBS compared to the Deg 1 RBS, the difference in eGFP production between these two strains was only a factor of 1.3. A similar discrepancy between predicted T7RNAP expression and actual eGFP production rate can be observed for the Deg 2 RBS, indicating that there is a surplus of T7RNAP in the original system. For the weaker RBS, it becomes apparent that the growth rate benefits from the lower eGFP production rate, proving once more that the system is suffering from a problem associated with the T7RNAP-driven eGFP production. Interestingly, the strain with the medium RBS had a lower eGFP production rate than the most degenerated variant of the original T7RNAP RBS (Deg 4), even though it was referred to as a medium-strength RBS and had the same predicted UTR value as Deg 3. A possible explanation could be that the strength of the RBS is also determined by the coding sequence, which was yellow fluorescent protein in the original work^*30*^ and T7RNAP here.

To test whether the RBS of the T7RNAP transcript influences the controllability of the system, we determined the eGFP production rate and growth rate of the RBS library strains in the absence of the inducer 3-MB. The controllability, in this case, is defined as the eGFP production rate for strains in the induced state divided by the corresponding eGFP production rate in the non-induced state. The controllability for all RBS library strains was in the range of 4±1, meaning that the RBS strength upstream of the *T7NRAP* gene did not influence the controllability of the system.

As a next step, we set out to test the RBS variants that performed best in combination with the construct containing the tight tZ terminator. This could be done by simply exchanging the plasmid-based module of the system. As shown in Figure 7, the saturation of the system with T7RNAP seems to be even more apparent within the tZ system, since even with the 3^rd^ degenerated RBS, the eGFP production rate was only slightly lower than with the original RBS (17748 ± 544.9 RLU OD_600_^-1^ h^-1^ vs. 22325 ± 1789 RLU OD_600_^-1^ h^-1^ respectively). This is in contrast to the original system, where the eGFP production rate in *P. putida* T7RNAP Deg 3 (pT7-eGFP) was 2.4 fold lower than in *P. putida* T7RNAP (pT7-eGFP). A possible explanation for this observation is that, due to the improved TE in the strain harboring the tZ terminator downstream of the *eGFP* gene, the expression of eGFP is more efficient, and therefore a certain threshold of eGFP is reached faster in the tZ-based system than in the system based on the TΦ terminator. Interestingly, the growth rate of *P. putida* T7RNAP Deg 3 (pT7-eGFP-tZ) is considerably higher than that of *P. putida* T7RNAP (pT7-eGFP-tZ), even though its eGFP production rate is only marginally lower, providing it with a beneficial trade-off between growth and production.

**Figure 7.**
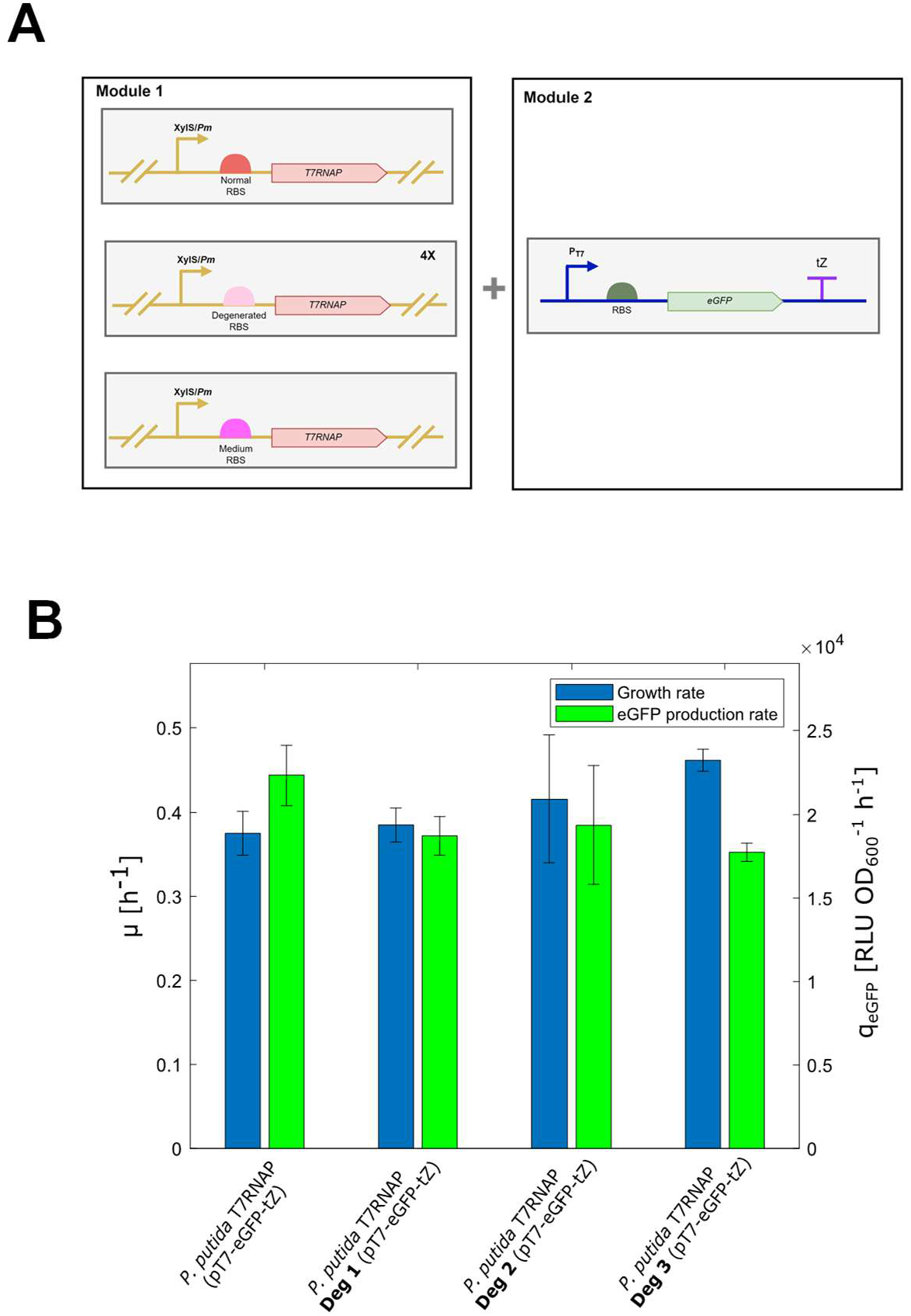
Testing the RBS library in combination with the pT7-eGFP-tZ plasmid. **A:** Genetic layout of the RBS library strains that contain the pT7-eGFP-tZ plasmid as the second module. Module 1 comprised the *T7RNAP* gene integrated into the genome, module 2 comprises a plasmid for T7RNAP driven eGFP expression. In total, four strains were tested. The difference between the strains was the RBS upstream of the *T7RNAP* gene in module 1. The predicted strength of the RBS sequences is gradually decreasing (compare table 1). Module 2 had the same composition for all strains and contained the strong synthetic terminator tZ downstream of the *eGFP* gene. **B:** Growth rate and eGFP production rate. Cultures were grown in 10 mL M9 Minimal medium with 3 g L-1 glucose as a C-source, at a temperature of 30°C and an agitation speed of 220 rpm. The cultures were induced with 0.1 mM 3-MB at the start of cultivation. The grey shaded area in B represents the results of non-induced strains as control. All cultivations were performed in triplicates. Error bars represent the standard deviation (based on a sample) of the replicates, calculated using the n-1 method.

### Theoretical considerations – kinetic analysis

After experimental confirmation of the T7RNAP saturation hypothesis, we focused on finding a kinetic expression to integrate the experimental data into a quantitative description. To stimulate the system, the synthesis of the T7RNAP was altered by changing the transcription initiation (via inducer amount) or translation initiation (via RBS sequence). The input factors here are the amount of 3-MB and the UTR expression values for the RBS upstream of the *T7RNAP* gene. Although the polymerase amount or activity was not measured directly, the fluorescence of eGFP and the eGFP production rate could be used as informative signals (outputs). The proposed model describes the influence of inducer and T7RNAP RBS sequence on the eGFP production rate in the form of Michaelis-Menten kinetics.

Starting from a mass balance equation for a single intracellular component *M*, an ordinary differential equation for the intracellular concentration *c*_*M*_ of the component reads as follows:

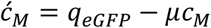

*q* is the sum of all synthesis rates (in our case, only a single process leads to eGFP synthesis), and µ is the specific growth rate. During exponential growth, a balance between synthesis and decay by dilution allows us to determine the rate of synthesis:

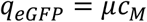

Given the experimental data of the fluorescence, representing the intracellular concentration of eGFP, and the specific growth rate, the rate of synthesis of eGFP for different strain variants was determined. To relate the production rate with the input factors (inducer concentration and UTR values), a Michaelis-Menten equation with two kinetic parameters is used:

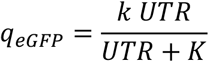

In the equation, *k* represents the maximal value of the rate, and *K* describes the concentration (in our case UTR value) that results in *k*/2 (half-saturation parameter). We used the following procedure to determine these parameters; parameter *k* is set to the maximal observed eGFP production rate for the strain with the strongest RBS upstream of the *T7RNAP* gene (see Table 1). In the second step, *K* is estimated by a simple regression using MATLAB. We used this procedure to determine parameters for the RBS library of the strain *P. putida* T7RNAP (pT7-eGFP) (compare Figure 6), for both the non-induced and the induced states. This results in two *k* values and only one single *K* value (the *K* value is the same for the induced and non-induced cases).

Starting with these parameters, we determined how the parameters adapt in the case of a stronger RBS upstream of the *eGFP* gene (*P. putida* T7RNAP (pT7-eGFP-strongRBS)) and for the modified termination sequence downstream of the *eGFP* gene (*P. putida* T7RNAP (pT7-eGFP-tZ)). These changes are expressed in two factors, *α* and *β*, that show an influence on *k* and *K*, respectively, with the formula:

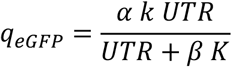

The results for the modified termination sequence (tZ) and the strong RBS upstream of the *eGFP* gene are as follows:

To summarize the results, the exchange of the termination sequence is described with constant factors in the kinetics for both the induced and the non-induced state. The kinetics allows us to predict the rate of synthesis of the protein if the UTR values are changed (Figure 8). Parameter *k* in the kinetics is a summation parameter taking into account transcription as well as translation characteristics, like the overall number of RNA polymerases, the overall number of ribosomes, and the overall number of binding sites for these control components. Parameter *K* can be seen as a summation parameter for the binding affinities of the polymerase and the ribosomes to their respective binding sites.

**Figure 8.**
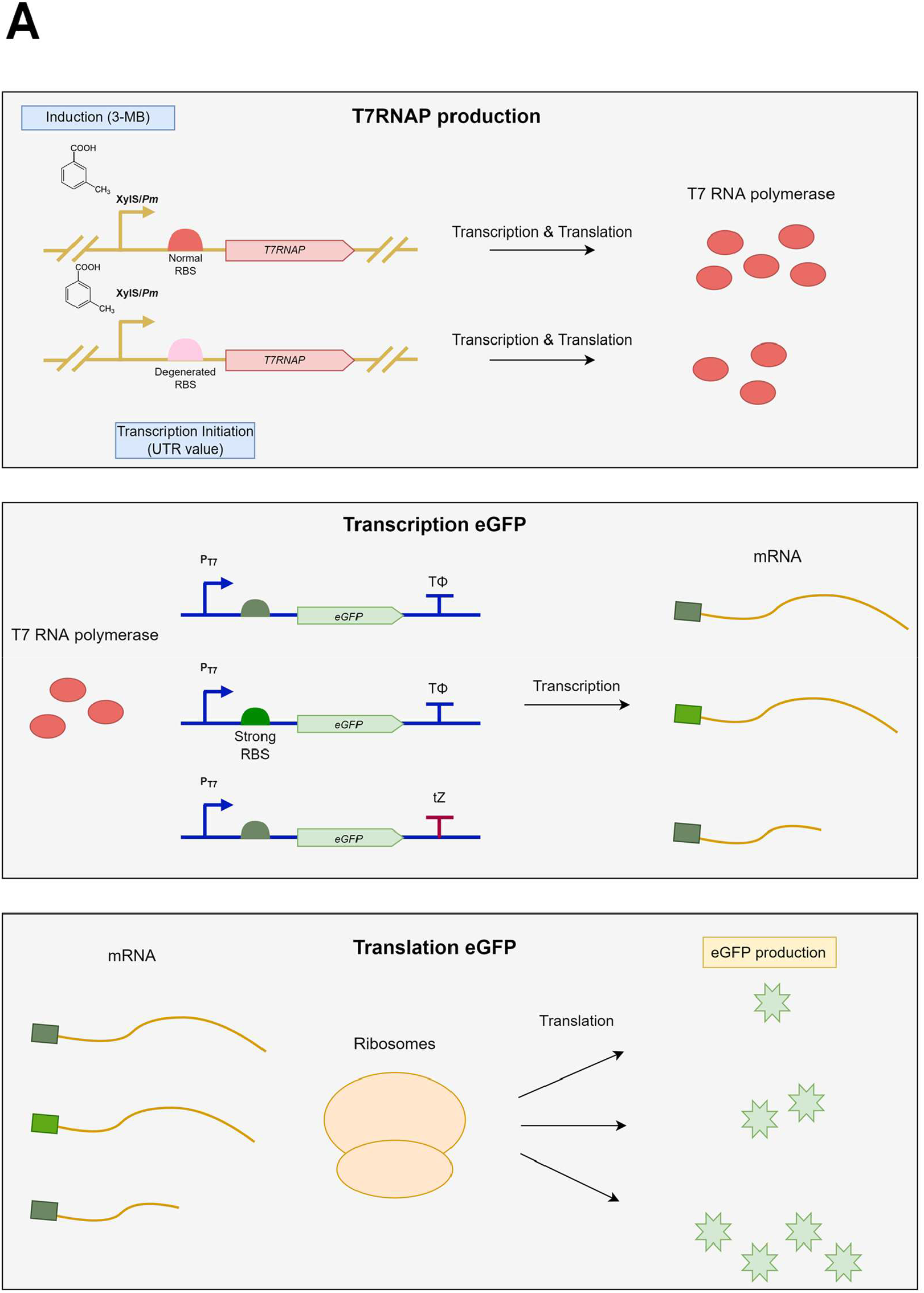
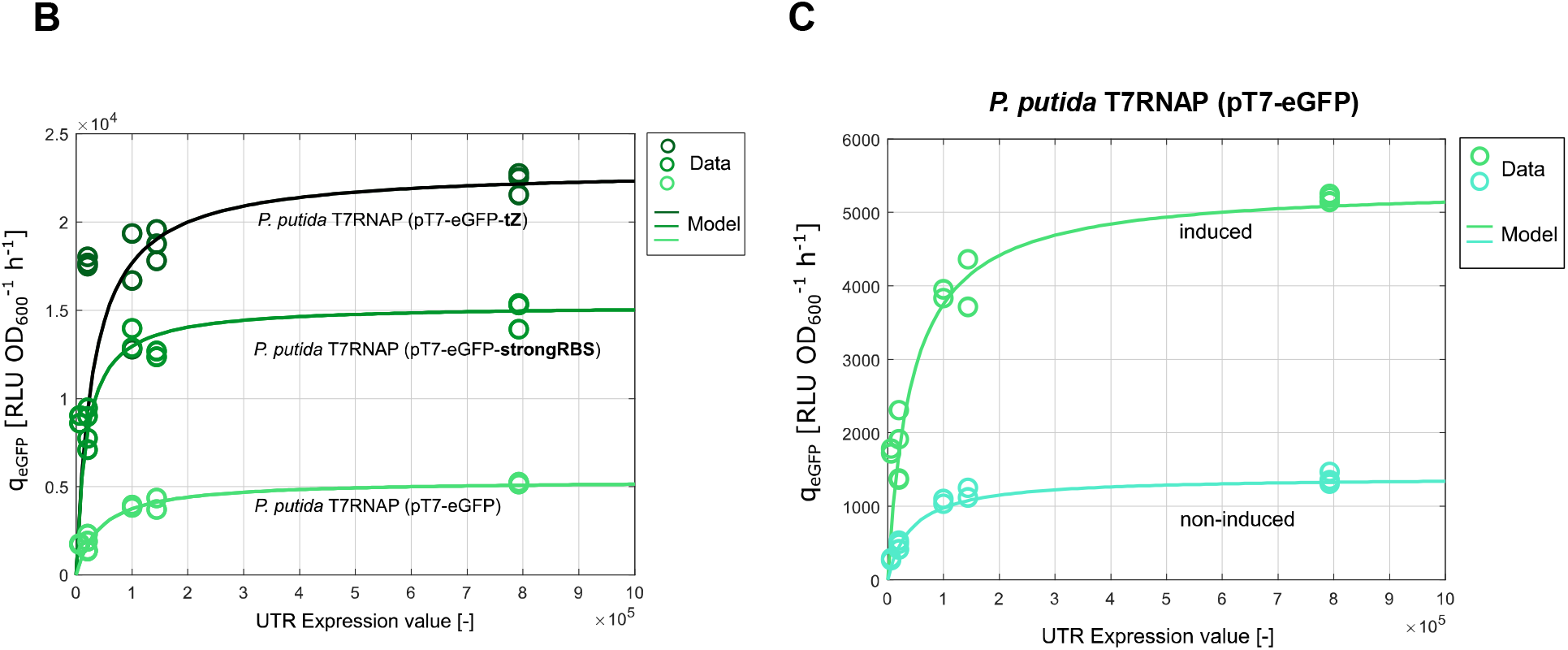
Representation of the kinetic model which describes the experimental data of the T7RNAP RBS libraries. **A:** Graphical description of the processes involved in the synthesis of eGFP with the T7RNAP RBS library. The first step is the synthesis of varying amounts of T7RNAP, depending on both, the RBS upstream of the *T7RNAP* gene and on the inducer concentration (blue boxes). Afterwards, different amounts of eGFP mRNA are synthesized, depending on the T7RNAP concentration. The mRNA length and composition depends on the architecture of the pT7-eGFP plasmid of the strain. The last step is the translation of the eGFP mRNAs by the ribosomes, where the rate of synthesis, the output (yellow box), depends on both the amount of available eGFP mRNA as well as the mRNA length and composition. **B:** Experimentally determined eGFP production rates for the different RBS sequences upstream of the *T7RNAP* gene (circles) and eGFP production rates as predicted by the kinetic model (curves), for three different plasmid architectures, in the case of full induction with 0.1 mM 3-MB. **C:** Experimentally determined eGFP production rates for the different RBS sequences upstream of the *T7RNAP* gene (circles) and eGFP production rates as predicted by the kinetic model (curves), for *P. putida* T7RNAP (pT7-eGFP), in the case of full induction (0.1 mM 3-MB) and in the absence of inducer.

Parameters *α* and *β* are the same for *P. putida* T7RNAP (pT7-eGFP-tZ), both in induced and non-induced states. This suggests that a cellular process other than the T7 RNA polymerase synthesis is responsible for the altered kinetics of this strain. For *P. putida* T7RNAP (pT7-eGFP-strongRBS), a nearly threefold increased value of *k* is observed, together with a smaller binding parameter *K*, confirming that the ribosomes’ binding of the mRNA transcript is truly stronger.

## Conclusion

Overall, a modular expression platform for T7RNAP-driven expression of heterologous genes in *P. putida* could be established and characterized. By reshaping the genetic circuit via replacing the TΦ terminator downstream of the *eGFP* gene and adapting the RBS in front of the *T7RNAP* gene, we efficiently tackled the problems associated with the original system. The application of the synthetic tZ terminator effectively prevented read-through transcription by the T7RNAP, increasing the eGFP production rate and growth rate of the strains. By varying the amount of T7RNAP in the system by changing the RBS upstream of the *T7RNAP* gene, the hypothesis of T7RNAP saturation was confirmed. In this context, improving the transcription initiation rate could potentially help counteract T7RNAP saturation, elevate mRNA levels, and increase protein production. Recent research by Shilling et al. already suggests that the T7 promotor as presented in the conventional pET28a vector could be optimized to enhance expression results.^*32*^ However, another possible explanation for the productive limitation of the T7RNAP system might be that translation becomes the rate-limiting step. Hence, further optimization could be more likely achieved by focussing on ribosome/transcript interaction rather than polymerase/gene interactions. To improve the controllability of the system, traditional approaches used for T7RNAP-based *E. coli* systems could be tested, e.g. T7RNAP lysozyme expression or applying transcription suppression as is the case with lacI/*T7* promotor.^*33, 34*^ Key to the optimization of the system is the choice of an optimal RBS for the T7RNAP. Ideally, it should be chosen in such a way that the system operates at the verge of saturation so that maximum eGFP production rate and growth are achieved without wasting cellular resources on surplus T7RNAP production. The presented model assists in this choice of T7RNAP, by indicating the relative strength of the RBS, taking into account the experimentally verified saturation effects.

## Materials & Methods

### Construction of plasmids

All constructs assembled in this work were essentially created using the plasmids pSEVA438 and pTn7-M^*35*^ as templates. Additional genomic parts / DNA fragments were generated by performing PCRs and incorporated into the various plasmids by performing restriction digestion and T4 ligation. Primers were supplied by Eurofins GmbH (Germany) or IDT (Leuven). Enzymes were ordered from New England Biolabs GmbH (Germany). Purification and extraction steps were performed using various kits from NIPPON Genetics EUROPE GmbH (Germany). A list of plasmids and primers used in this work can be found in the Supporting Information (Table S2 and S3).

### Transformation and conjugation

Using the method of Chung et al., newly assembled vectors were transformed into competent *E. coli* DH5α λ-*pir* cells.^*36*^ After growth on LB-Agar with appropriate antibiotics, the resulting colonies were checked for the new construct by performing colony PCR. After confirmation by Colony PCR, plasmids were sent to Eurofins GmbH (Germany) or Genewiz (Germany) for sequencing. Plasmids were transferred to *P. putida* either by triparental (in case of pSEVA plasmids) or quatroparental (in case of pTn7-mini plasmids) conjugation using helper strains (see Supporting Information S1/Table S1).^*37*^

### Cultivation, medium and sampling

Pre-cultures were grown overnight in Lysogeny Broth (LB) medium supplemented with either 200 µg/mL streptomycin or 10 µg/mL gentamycin, depending on the antibiotic resistance of the strain. Main cultures were grown in 100 mL shake flasks with a medium volume of 10 mL and were inoculated with either 100 µL or 200 µL of overnight LB pre-culture. The medium used for cultivation was M9 minimal medium^*38*^ with 3 g/L glucose as a carbon source. In the case of induction of cultures, 0.1 mM 3-MB was added at the start of the cultivation. The cultures were incubated at 30°C in a rotary shaker (Thermofisher, USA) at an agitation speed of 220 rpm. Approximately every hour, 25-200 µL (depending on dilution factor) of sample were taken from the cultures and optical density (600 nm) and eGFP fluorescence (at 480Ex_515Em with gain 80 and at 480Ex_530Em with gain 60) were measured in a microplate reader (TECAN, Switzerland). Absorbance and fluorescence of the samples were corrected using Phosphate Buffer Saline (PBS) as a blank.

### eGFP quantification

In this work, eGFP is expressed in relative luminescence units (RLU). To determine the RLU, a procedure similar to the one described by Lichten et al. was followed.^*39*^ Therefore, both specific and unspecific eGFP fluorescence signals were regarded. Additionally, the interfering fluorescence of cell components has to be considered, and the specific (480Ex_515Em) and unspecific (480Ex_530Em) signals must be corrected for this term. The actual measured specific (F_1_) and unspecific fluorescence (F_2_) signals are composed as followed:

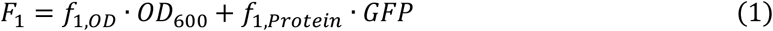

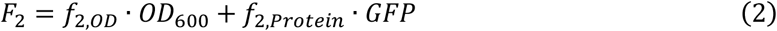

Where f_OD_ indicates the portion of fluorescence caused by cell component fluorescence and f_Protein_ indicates the portion of fluorescence caused by the actual presence of the eGFP protein. When equation 1 and 2 are divided by their respective f_OD_ and equation 2 is subsequently extracted from equation 1, the following expression is obtained:

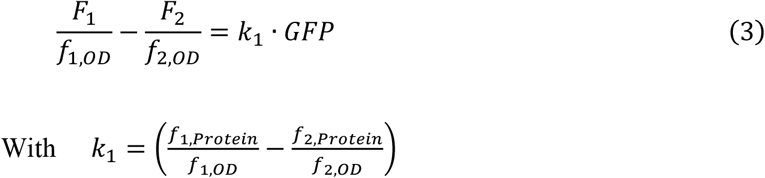

Multiplication of expression 3 with f_1,OD_ yield the following final expression:

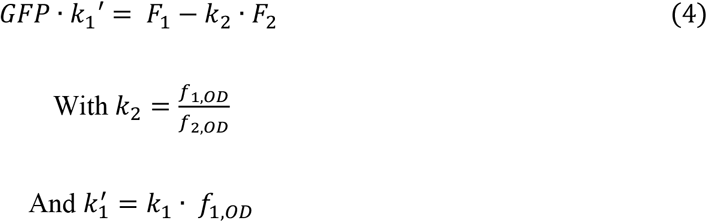

Factor k_2_ can be determined experimentally by correlating F_1_ and F_2_ for a non-GFP-producing strain. Using this correlation, eGFP can be semi-quantified. Since k_1_’ remains constant, the left-handed side of equation 4 correlates only with eGFP concentration and represents the RLU. The RLU changes over time and was calculated for each replicate at each sampling point.

To calculate the eGFP Yield (Y_eGFP_) a linear regression was performed. Since yield does not provide any information about the dynamics of GFP production, the GFP production rate (q_eGFP_) was calculated:

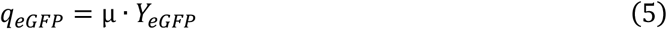

### *In vitro* T7RNAP transcription assay

The *in vitro* T7 transcription was performed according to the User Guide: Conventional *in vitro* Transcription of Thermo Fisher Scientific (USA). Shortly, a transcription puffer (Thermo Fischer Scientific (USA)), 2 mM of rNTPs (New England Biolabs GmbH (Germany)), template DNA, 30 units of T7RNAP (Thermo Fischer Scientific (USA)), and nuclease-free water were mixed. Then, the assays were incubated at 37°C. Assays were mixed in a sterile environment to avoid contamination with RNases. After two hours, the reaction was stopped by adding 2 µL 0.5 M EDTA (pH 8.0) and incubation at 65°C for 10 min. After the assay was completed, the RNA samples were immediately stored on ice, to prevent degradation of mRNA.

A negative control was included to identify whether the bands on the RNA agarose gel originated from mRNA rather than the template DNA. For this purpose, an assay was performed where the T7RNAP was replaced by deionized water.

To visualize the RNA synthesized in the T7 Transcription assay, a denaturing, 1% agarose, gel was cast. First, the agarose was mixed and dissolved in deionized water by heating in the microwave. After cooling the mixture to 60°C, DNA stain, MOPS Buffer (1x) and 37% formaldehyde (18 % (v/v)) were added. Thereafter, the gel was cast in the same manner as a normal DNA agarose gel.

In the next step, the samples were mixed with a loading dye. The loading dye was prepared by mixing 10 µL formamide, 4 µL formaldehyde (18%), 2 µL MOPS Buffer (10x), and 1 µL NEB Purple Gel Loading Dye (6x). To ca. 10 µL of the sample, 17µL of loading dye was added. The dyed samples were then incubated at 67°C for 5 minutes, to denature the mRNA. Afterward, the same procedure for loading, running, and visualizing was followed as described for a normal DNA agarose gel.

## Supporting information

Supporting Information (strains,plasmid,oligonucleotides)

## Associated content

### Supporting information

List of strains, plasmids and oligonucleotides used during this work.

## Author Information

### Author contribution

H.L. and K.P.-G. conceptualized the project, M.B. was responsible for the investigation. H.L., K.P.-G. and M.B. were involved in analysis and interpretation of the data. M.B. wrote the experimental part of the manuscript, A.K. performed the theoretical analysis of the data and wrote the theoretical part of the manuscript. H.L. and K.P.-G. were involved in the review and editing of the manuscript. A.K. was responsible for project administration and funding acquisition. All authors read and approved the final manuscript.

## Acknowledgments

We would like to thank Victor de Lorenzo (CNB-CSIC, Madrid) for providing the plasmid pSEVA438 and Gerald Striedner (BOKU, Vienna) for providing the tZ terminator. We would also like to thank Ana Sofia Ortega (TU Munich) for providing the strain *P. putida* (pSEVA438-eGFP) and for the experimental support. This research did not receive any specific grant from funding agencies in the public, commercial, or not-for-profit sectors.

