## Supporting Information (strains,plasmid,oligonucleotides) for "A modular T7-based gene expression platform in *Pseudomonas putida*: Construction and *in silico* analysis"

for

### Strains

S1. List of bacterial strains used in this work

| Strain | Description | Reference |
|---|---|---|
| *E. coli* HB101 (prk600) | Helper for triparental conjugation | Sambrook et al., 1989[1] |
| *E. coli* DH5α (pTns1) | Strain encoding the Tn7 Transposase. Required for genomic integration of pTn7 based plasmids upon conjugation | Choi et al., 2005[2] |
| *E. coli* DH5α λ pir | Host for recombinant transformation (containing λ pir lysogen from DH5α) | Kolter et al., 1978[3] |
| *P. putida* KT2440 | Safe strain gram-negative soil bacterium used for cloning and gene expression. Derived from *P.putida* mt-2 | Bagdasarian et al., 1981[4] |

### Plasmids

S2. List of available and newly created plasmids used in this work

| Plasmid | Size | Description | Reference |
|---|---|---|---|
| pSEVA438 | 5140 bp | Plasmid designed according to Standard European Vector Architecture. Antibiotic resistance: Sm/Sp<br>ORI: pBBR1<br>Cargo: XylS/*Pm* (Inducible with *3*-MB) | Silva-Roche et al., 2013[5] |
| pT7-eGFP | 4041 bp | pSEVA438 based vector. The XylS/*Pm* cassette is exchanged with eGFP under the control of the T7RNAP specific promoter, derived from the pET21b(+) plasmid | pET (Studier and Moffat, 1986)[6]<br>Plasmid (This work) |
| pT7-eGFP-ΔRBS | 4027 bp | pT7-eGFP containing a deletion of RBS upstream of the *eGFP* gene | This work |
| pT7-eGFP -strongRBS | 4059 bp | pT7-eGFP, where the original RBS upstream of the *eGFP* gene is exchanged with a strong synthetic RBS | Strong RBS (Ceroni et al., 2015)[7]<br>Plasmid (This work) |
| pET-kT7TZ | 5421 bp | pET30a vector containing the T7 promotor followed by the TZenit (tZ) Terminator (consisting of T3 -, rrnBT1- and TΦ terminator) | (Mairhofer et al., 2015)[8] |

| | | | |
|---|---|---|---|
| pT7-eGFP-tZ | approx. 4140 bp | pT7-eGFP where the inherent T7 terminator TΦ is replaced by the designed T7 terminator (tZ) | tZ terminator (Mairhofer et al., 2015)[8]<br>Plasmid (This work) |
| pTn7-M-XylS/Pm-T7-Deg (4x) | approx. 8471 bp | Mini-Tn7 plasmid for genomic integration of the XylS/*Pm* expression cassette followed by the *T7RNAP* gene with 4 gradually increasing degradations of its RBS | This work |
| pTn7-M-XylS/Pm-T7-MediumRBS | approx. 8471 bp | Mini-Tn7 plasmid for genomic integration of the XylS/*Pm* expression cassette followed by the *T7RNAP* gene with a medium strength synthetic RBS | Medium RBS (Levin-Karp et al., 2013)[9]<br>Plasmid (This work) |

### Oligos

S3. Sequence and description of oligonucleotides used for this work. In primer for cloning, the enzyme restriction site is **bolted**, and the annealing part of the primer is underlined.

| Name | Sequence (5'→ 3') | Description |
|---|---|---|
| Fwd_Primer_ RBS_Deg1 | ATAT**GAATTC**ATTTACTAACTGGAAGAGACACTAAATGAACACGATTAACATCGCTAAG | Amplification of T7RNAP containing 1st RBS Degradation (EcoRI restriction site) |
| Fwd_Primer_ RBS_Deg2 | ATAT**GAATTC**ATTTACTAACTGTAAGAGACACTAAATGAACACGATTAACATCGCTAAG | Amplification of T7RNAP containing 2nd RBS Degradation (EcoRI restriction site) |
| Fwd_Primer_RBS_Deg3 | ATAT**GAATTC**ATTTACTAACTGTAATAGACACTAAATGAACACGATTAACATCGCTAAG | Amplification of T7RNAP containing 3rd RBS Degradation (EcoRI restriction site) |
| Fwd_Primer_RBS_Deg4 | ATAT**GAATTC**ATTTACTAACTGTAATCGACACTAAATGAACACGATTAACATCGCTAAG | Amplification of T7RNAP containing 4rd RBS Degradation (EcoRI restriction site) |
| Fwd_Primer_RBS_Milo | ATAT**GAATTC**TTCGCAGGGGGAAGATGAACACGATTAACATCGCTAAGAACG | Amplification of T7RNAP containing medium strength RBS (EcoRI restriction site) |
| Rev_Primer_RBS_Deg | TAAT**AAGCT**TGCATGCCTGCAGTTACGCGAACGCGAAGTC | Amplification of T7RNAP, reverse Primer (for cloning Deg1-4 and Medium RBS) (HindIII restriction site) |
| Fwd_Primer_RBS_Ohne | ATAT**CTGCA**GATGGTGAGCAAGGGCGAG | Amplification of eGFP without RBS (PstI restriction site) |
| Fwd_Primer_RBS_Ceroni | ATTA**CTGCA**GTACTAGAGAAATCAAATTAAGGAGGTAAGATAATGGTGAGCAAGGGCGAG | Amplification of eGFP with stronger RBS (PstI restriction site) |
| Fwd_Primer_tZTerminator | ATAT**ACTAGT**CCGTCGACCTAGCATAACC | Amplification of Terminator tZ (SpeI restriction site) |
| Rev_Primer_tZTerminator | AGCAA**TTTAAAT**GGATATAGTTCCTCCTTTCAGC | Amplification of Terminator tZ (SwaI restriction site) |

| | | |
|---|---|---|
| Fwd_ Sequenzierung_tZ | GGATAACAATTCCCTCCTAGG | Forward primer used for verifying the cloning results of the tZ terminator and for sequencing of pT7-eGFP (pET) plasmid |
| Rev_Sequenzierung_tZ | CCGAGGCATAGACTGTACC | Reverse primer used for verifying the cloning results of the tZ terminator |
| Fwd_238 | TATCTCTAGTAAGGCCTACC | Forward primer for verifying cloning results in pSEVA438 and pTn7-XylS/Pm-T7 |
| Rev_224 | ATCCAGATGGAGTTCTGAGG | Reverse primer for verifying cloning results in pSEVA438, pT7-eGFP and pTn7-XylS/Pm-T7 |
| SW_16 | GCCTTTCGTTTTATTTGATGCCT | Forward primer for verifying cloning results in pT7-eGFP and for generating ReadThrough DNA sequences |
| $P_{put\text{-}glmSUP}$ | AGTCAGAGTTACGGAATTGTAGG | Forward primer for verifying correct genomic integration of the pTn7 mini plasmid [2] |
| $P_{Tn7L}$ | ATTAGCTTACGACGCTACACCC | Reverse primer for verifying correct genomic integration of the pTn7 mini plasmid [2] |
| ReadThrough_1 | AAGGTCGTTGATCAAAGCTCG | Anneals ca 500 bp downstream of TΦ terminator in pT7-eGFP |
| ReadThrough_2 | CAAGCGATCTTCTTCTTGTCC | Anneals ca 1000 bp downstream of TΦ terminator in pT7-eGFP |
| ReadThrough_3 | GAAACGGAGGAATGGGAACG | Anneals ca 1500 bp downstream of TΦ terminator in pT7-eGFP |
| ReadThrough_4 | GAGAAATCGGCATTCAAGCC | Anneals ca 2000 bp downstream of TΦ terminator in pT7-eGFP |
| ReadThrough_5 | AATTTTCTCTGGGGAAAAGCC | Anneals ca 2500 bp downstream of TΦ terminator in pT7-eGFP |
